# Cervical spinal cord stimulation disrupts proprioception yet improves voluntary arm reaching

**DOI:** 10.64898/2026.06.20.733548

**Authors:** Erick Carranza, Roberto de Freitas, Nikhil Verma, Luigi Borda, Erynn Sorensen, Amy Boos, George Wittenberg, Lee Fisher, Marc P. Powell, Peter Gerszten, Douglas Weber, John W. Krakauer, Jonathan Tsay, Marco Capogrosso, Elvira Pirondini

## Abstract

Whether proprioception is necessary for upper-limb motor control has been debated for decades. Classic studies in deafferented animals and humans suggested that proprioception may be dispensable for rapid, goal-directed movements. However, chronic sensory loss conflates the absence of proprioceptive input with years of compensatory adaptation. As a result, the field has lacked a strong causal test of proprioception’s contribution to motor control. Here, we leveraged a clinical trial of cervical spinal cord stimulation (SCS) in individuals with chronic post-stroke hemiparesis to study if electrical stimulation of sensory afferents causally perturbs proprioception and affects arm reaching. We found that turning SCS ON causally impaired proprioceptive perception and postural stabilization in response to force perturbations, and enhanced adaptation to visual errors during implicit learning. Yet visually and non-visually rapid, goal-directed reaching improved in smoothness, straightness and spatial accuracy. These findings provide strong causal evidence that proprioception is not required for rapid, goal-directed action, helping resolve a decades-long debate regarding its necessity for effective movement.

## Introduction

A long line of work in non-human primates suggests that proprioception—the sense of limb position and motion—is not strictly required to produce fast, goal-directed reaches^1–3^. Specifically, large fiber deafferentation or dorsal column lesions leave the ability to reach toward targets largely intact, albeit with reduced accuracy and increased reliance on vision. Converging evidence comes from neuropsychological studies of deafferented patients, who, despite profound sensory loss due to peripheral neuropathy, can execute relatively accurate ballistic reaching movements^4,5^.

However, these findings provide only indirect evidence, as proprioception cannot be selectively and reversibly manipulated in deafferented patients or chronically lesioned animals. Specifically, chronic sensory loss conflates the absence of proprioception with years of compensation and learned reliance on vision and other sensorimotor signals^6–8^. As a result, it remains unclear whether rapid, goal-directed movements are truly independent of proprioception or whether the nervous system can somehow adapt to its long-term loss^9–12^. Yet, existing methods for perturbing proprioception in humans are limited: tendon vibration is unreliable and produces only partial, indirect disruption^13,14^, while transient perturbations such as cooling or ischemic nerve block are non-selective and offer limited temporal control^15,16^. Establishing causal manipulations in humans is therefore essential to determine whether proprioception is necessary for upper-limb motor control.

Recent work provides a promising path forward: leveraging clinical trials of epidural spinal cord stimulation (SCS)^17–30^. SCS selectively stimulates dorsal root sensory afferents that project to lower– and upper-limb spinal motor neuron pools in humans^31,32^. Through this excitatory monosynaptic pathway, SCS can increase motoneuron excitability facilitating strength and limb motor performance^19,23,28^. Critically, SCS primarily recruits large-diameter proprioceptive afferents (group Ia from muscle spindles and Ib afferents from Golgi tendon organs^31,33^). However, because this activation is tonic and uncorrelated with limb movement, it is thought to mask the natural, movement-evoked afferent discharge that conveys proprioceptive information, providing a means to causally perturb proprioceptive signals^34^. Indeed, continuous lumbar SCS has been shown to improve lower-limb motor performance^23,35,36^, while simultaneously impairing knee proprioceptive acuity in individuals with spinal cord injury^34^.

Here, we leveraged a clinical trial of cervical SCS in people with chronic post-stroke hemiparesis (clinical trial: NCT04512690^19,29^) to understand if and how electrical stimulation of the cervical proprioceptive afferents impairs proprioception acuity in humans and how these impairments influenced motor control. To do so, we designed a comprehensive battery of sensorimotor tasks, including joint position sense, kinesthesia (the perception of limb motion, assessed here via detection threshold for passive displacement), feedback responses to mechanical perturbations of the arm, and implicit visuomotor adaptation. We found that, despite clear degradation in proprioceptive perception, postural stabilization in response to mechanical perturbations, and visuomotor adaptation, rapid, goal-directed reaching improved with stimulation even without visual feedback. Together, these results provide causal evidence that proprioception is not required for rapid, goal-directed actions and establish cervical SCS as a tool for interrogating the role of proprioception in human behavior.

## Results

### Spinal cord stimulation directly targets spinal sensory afferents

Our experiments were executed in the context of an exploratory clinical trial (NCT04512690) testing the preliminary feasibility of SCS to improve upper-limb motor function in people with post-stroke hemiparesis^19,29^. Participants (Supplementary Table 1) were implanted with two percutaneous 8-contact epidural leads in the dorsal cervical spinal cord, targeting the C3 to T1 spinal segments (Figure 1a). Fluoroscopic imaging during surgery confirmed rostro-caudal coverage across the intended spinal segments (Figure 1b, Supplementary figure 1c). These segments were selected to enable stimulation of the sensory afferents innervating proximal and distal upper-limb muscles via pre-synaptic recruitment of spinal α-motoneurons from the sensory afferents (Figure 1c).

**Fig. 1.**
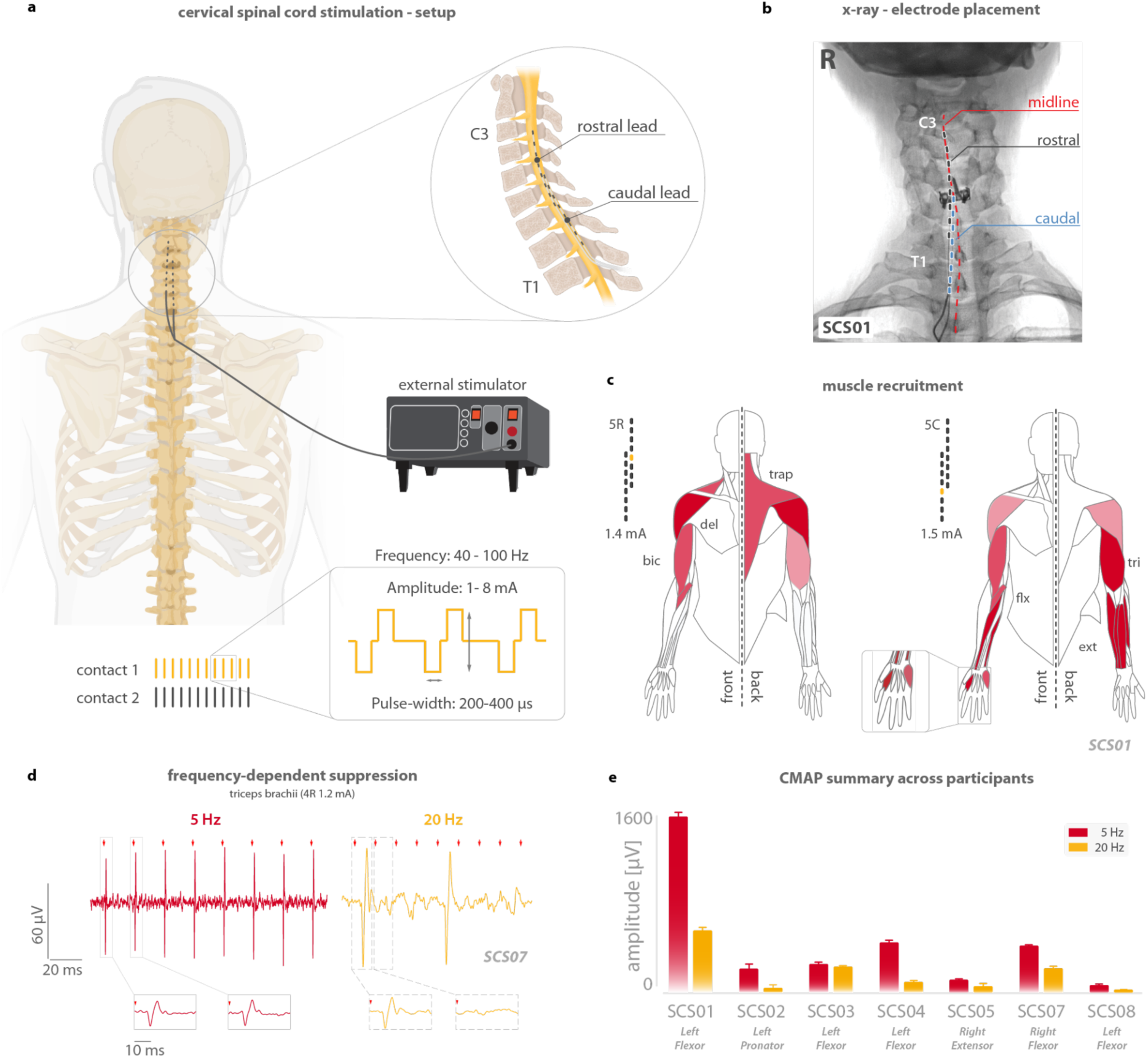
| Cervical spinal cord stimulation experimental setup and upper-limb muscle recruitment. **a.** Overview of the experimental setup. Participants were implanted with two 8-contact epidural leads spanning C3–T1 cervical segments and connected to an external pulse stimulator. Rostral–caudal lead placement targets segment-specific motor pools within the cervical spinal cord. Stimulation waveforms consisted of 200–400 µs biphasic pulses delivered at 1–6 mA, with frequencies ranging from 5–100 Hz. **b.** Intraoperative X-ray for participant SCS01 confirms bilateral epidural placement centered over the cervical spinal cord midline. **c.** Muscle recruitment map for SCS01 across stimulation amplitudes at rostral (5R) and caudal (5C) contacts, showing activation of proximal (e.g., deltoid, biceps) and distal (e.g., wrist extensors) upper-limb muscles. Muscle recruitment included both flexor and extensor groups, with side-specific selectivity depending on electrode location and current amplitude. Abbreviations: trap, trapezius; del, anterior deltoid; bic, biceps; tri, triceps; flx, flexor; ext, extensor. **d.** Frequency-dependent suppression of compound muscle action potentials (CMAPs) recorded in SCS07 during low (5 Hz, red) and high (20 Hz, yellow) frequency stimulation. Insets show representative CMAP traces. **e.** Across participants, CMAP amplitude was consistently reduced with stimulation at 20 Hz relative to stimulation at 5 Hz, indicating suppression of motor responses at higher frequencies and confirming frequency-dependent recruitment modulation. CMAP amplitude was quantified as the averaged peak-to-peak voltage of the three deflections following the stimulation artifact for each participant during intraoperative testing. Different muscles are shown across participants because intraoperative electrophysiological testing targeted the muscles that exhibited the most reliable stimulation evoked responses in each individual. Bars represent mean values and error bars indicate the standard error of the mean (SEM).

To confirm appropriate targeting of the dorsal root entry zones and trans-synaptic activation of spinal motor neurons, we assessed the presence of frequency-dependent suppression of compound muscle action potentials (CMAPs), i.e., the summed electrical response of a muscle following stimulation of its motor nerve (Figure 1d). In all participants, low-frequency (5 Hz) stimulation reliably evoked large CMAPs in upper-limb muscles, while high-frequency stimulation (20 Hz) led to progressively diminished responses, consistent with classical rate-dependent depression mediated by Ia afferent pathways^37,38^ (Figure 1e). This electrophysiological signature confirms pre-synaptic recruitment of spinal α-motoneurons via sensory afferents and not through direct activation of motor axons, which would not cause a decrease in responses at high-frequency stimulation.

During behavioral testing, stimulation was delivered continuously through one or more contacts^19,29^. For each participant, we identified stimulation configurations that promoted activity in the affected muscles (Figure 1c, see methods). As expected and previously reported^19,34,39^, all participants but SCS05 reported paresthesia over the arm during the stimulation sessions. We previously showed immediate changes in motor performance when stimulation was turned ON compared to stimulation OFF: these included changes in upper-limb strength, movement smoothness, and reduction of spasticity^19,29^. After 29 days, the leads were removed. All participants tolerated the procedures and stimulation sessions without severe adverse events (see de Freitas et al. 2026 for more details)^29^.

### Spinal cord stimulation selectively impairs kinesthesia but spares static position sense

Given that SCS preferentially recruits large-diameter sensory afferents, we tested whether stimulation disrupts proprioceptive processing. Specifically, we assessed the impact of SCS on two aspects of proprioception: kinesthesia, measured as the threshold for detecting passive limb motion, and static position sense, measured as the accuracy of matching a held limb position without vision.

We employed the threshold to detection of passive movement (TTDPM) task in the elbow joint, a validated measure of kinesthetic acuity^34^. A slow, robot-controlled elbow flexion or extension movement (Figure 2a) was applied randomly to the impaired arm, while participants were instructed to press a button and verbally report the direction of movement as soon as they perceived it (Figure 2b).

**Fig. 2.**
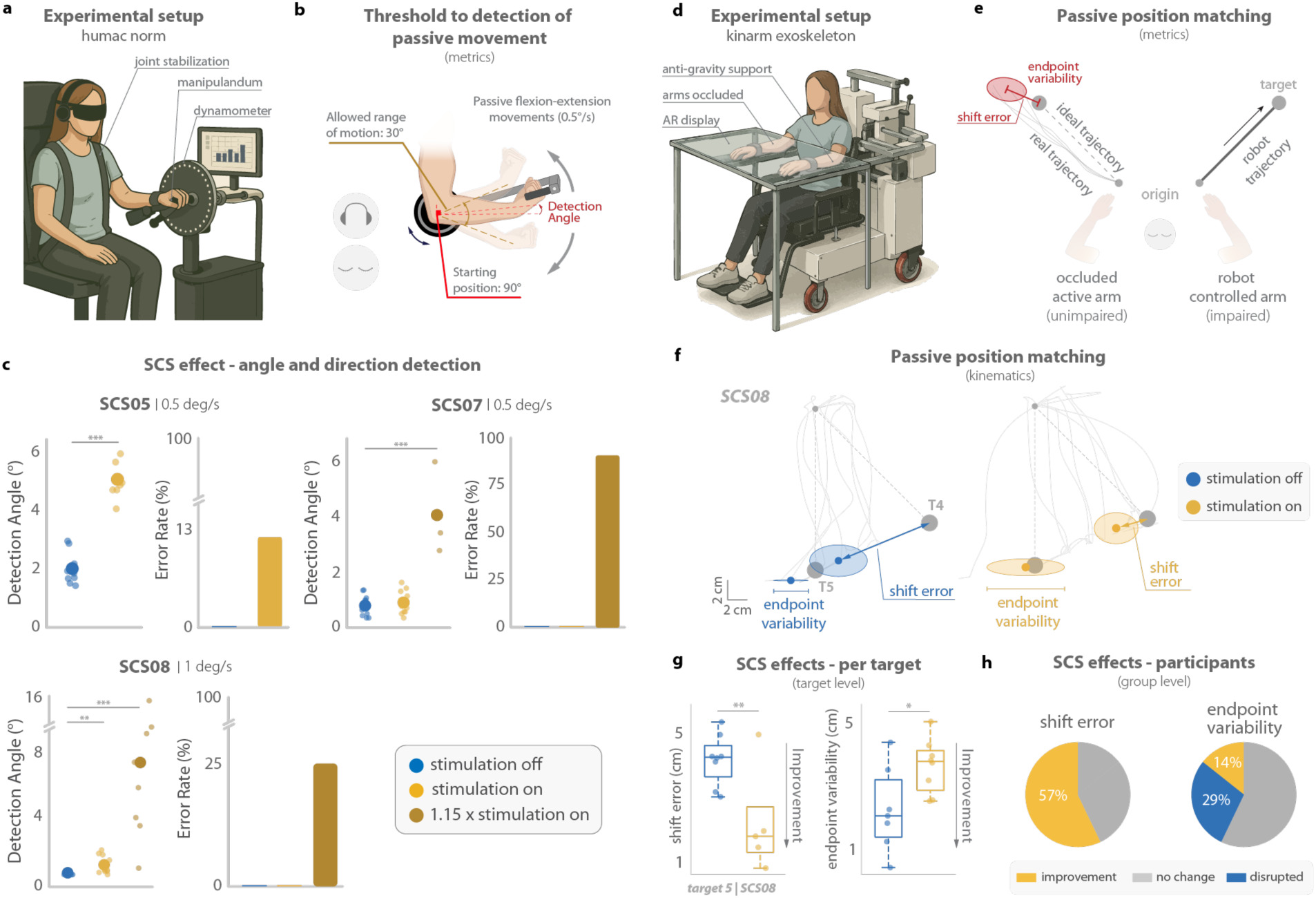
| Spinal cord stimulation disrupts dynamic proprioceptive processing but spares static position sense. **a.** Dynamic proprioceptive sense (kinesthesia) was measured using the threshold to detection of passive movement (TTDPM) task on the isokinetic dynamometer (HUMAC Norm). **b.** Participants reported the onset and direction of robot-driven flexion-extension movements applied to the impaired arm while vision was occluded by a blindfold and audition was masked by white noise. **c.** TTDPM results from three participants (SCS05, SCS07, SCS08). Stimulation increased the detection angle as well as the direction error rates, consistent with impaired kinesthesia. Increasing stimulation amplitude (1.15x), amplified the disruptive effect both in detection angle and error rate. Large dots represent mean detection angles. Individual dots represent single trials. Blue indicates stimulation OFF, yellow indicates stimulation ON, and darker yellow indicates increased stimulation amplitude. **d.** Experimental setup for the passive position matching task (PPM) on the KINARM upper-limb exoskeleton. The robotic platform allows for gravity support and occlusion of the upper limbs. A virtual reality display provides visual feedback according to the task requirements. **e.** The robot passively displaced the impaired arm to a target position; participants mirror-matched this location with their unaffected arm without visual feedback. **f.** Example of PPM kinematic traces illustrating SCS effects on shift error (accuracy) and endpoint variability (precision) for one participant (SCS08). **g.** Per-target comparison of shift error and endpoint variability with and without stimulation (SCS08). Colors indicate stimulation condition (blue: stimulation OFF; yellow: stimulation ON). Improvement was defined according to the expected direction of better performance for each metric, where a reduction in shift error and endpoint variability indicate improvement. Box plots represent the median, 25th and 75th percentiles and minimum and maximum data points. Individual dots represent single trials. **h.** Summary of stimulation effects on proprioceptive accuracy (shift error) and precision (variability) across participants. Pie charts show the proportion of participants who improved (yellow), worsened (blue), or showed no change (grey) under stimulation. Statistical significance was assessed with two-sided bootstrapping (n = 10 000): ^∗^*p* < 0.05, ^∗∗^*p* < 0.01, ^∗∗∗^*p* < 0.001.

In the absence of SCS, participants consistently detected arm motion and correctly identified the direction of displacement in every trial (Figure 2c). Strikingly, turning SCS ON produced an immediate increase in detection thresholds across all participants, indicating reduced kinesthetic sensitivity. Thresholds rose by 141%, 11%, and 53% for SCS05, SCS07, and SCS08 respectively, with SCS05 also exhibiting direction misclassifications (an increase from 0% to 11% error rate).

At higher stimulation amplitudes, kinesthetic performance further deteriorated (Figure 2c): in SCS07, detection thresholds increased from 1.70° to 4.56° (95% CI: 3.63, 6.25; p = 0.001), with direction errors on 91% of trials. In SCS08, thresholds rose from 0.89° to 7.23° (95% CI: 4.58, 10.66; p = 0.001), with a 25% error rate. The marked increases in detection angle and in direction misclassification demonstrate that stimulation intensity directly degrades kinesthesia.

To assess static position sense, we used a validated position-matching task^40–43^ in which the robot passively displaced the impaired arm to a target location, and participants mirror-matched the perceived position with their unimpaired arm (Figure 2e). The task was performed without vision and without active movement of the stimulated limb, isolating proprioceptive processing based on static limb position.

Across our two primary dependent measures^41^—shift error (the distance between the true mirrored position and the participant’s match) and precision (endpoint variability, reflecting trial-to-trial dispersion)—there were no consistent differences between SCS ON and SCS OFF conditions across participants. Specifically, for trial-to-trial variability one participant showed improvement and two participants showed degraded performances when the stimulation was turned ON compared to stimulation OFF; while for shift error only four participants had lower values, i.e., better performances, with stimulation ON. Importantly, these differences in the SCS effects across participants did not correlate with sensory impairments as captured by Fugl-Meyer (see Supplementary Table 1). Moreover, these results were validated by comparable performances in the Fugl-Meyer Sensory Score, in which blindfolded participants are required to mirror-match limb position, between stimulation ON and OFF (see Supplementary Table 3).

In summary, these findings indicate that cervical SCS selectively impairs kinesthesia while sparing static position sense. More broadly, they support the view that dynamic and static components of proprioception are mediated by distinct afferent populations and likely engage different spinal pathways^43–45^.

### Spinal cord stimulation enhances sensorimotor adaptation to visual perturbations

Does SCS-induced disruption of proprioception propagate to downstream motor function? To test this, we examined its impact on sensorimotor adaptation—the process by which motor commands are implicitly recalibrated to compensate for changes in the body (e.g., muscle fatigue) or the environment (e.g., added load). We assessed adaptation using the clamped feedback paradigm^46,47^. In this task, participants are instructed to reach directly toward a visual target while ignoring a feedback cursor that follows a fixed angular trajectory (“clamp”) offset from the target, independent of their actual hand movement (Figure 3a). Despite these instructions, the clamped visual feedback reliably induces gradual shifts in heading angle away from the target—a hallmark of sensorimotor adaptation.

**Fig. 3.**
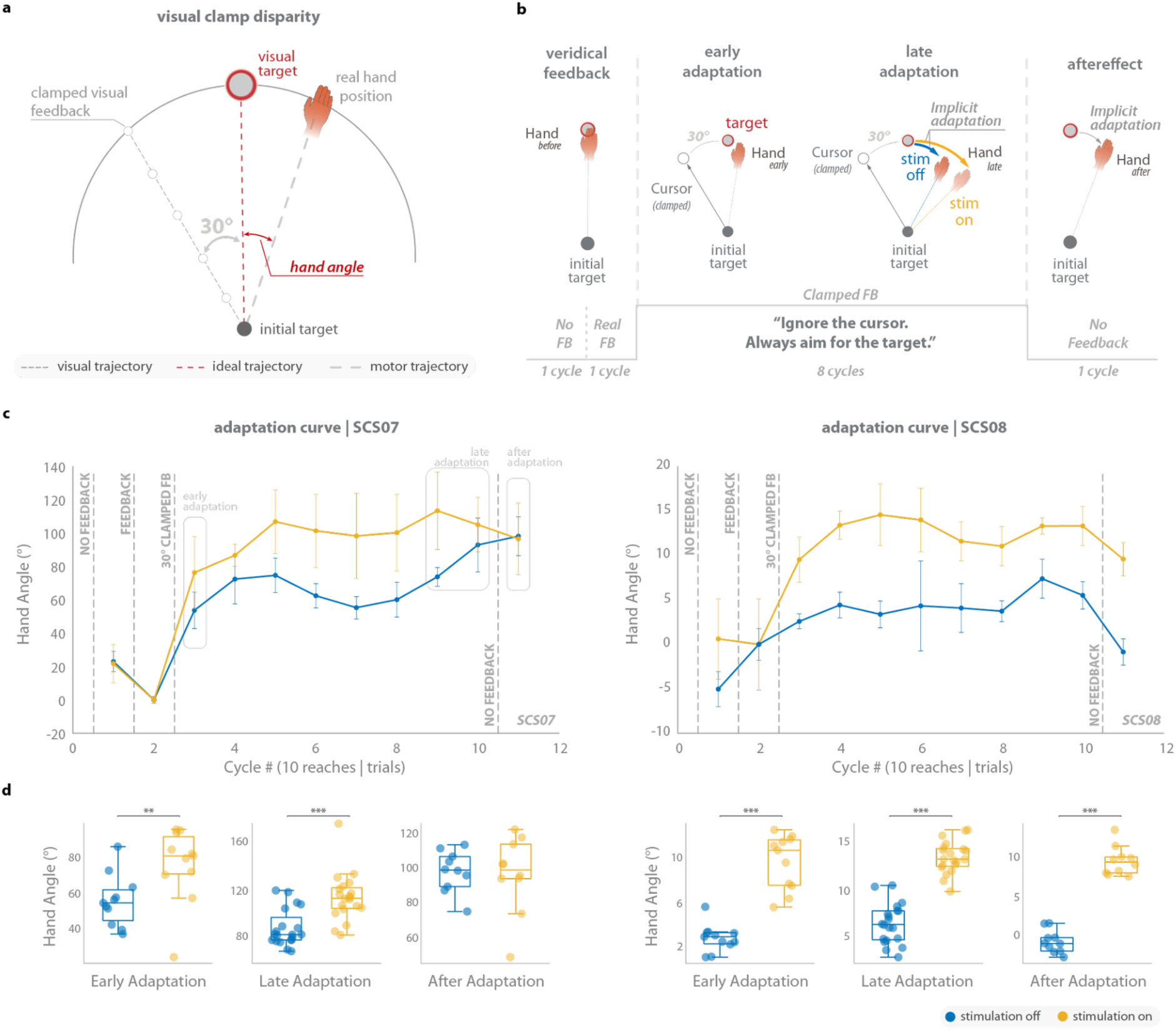
| Spinal cord stimulation enhances sensorimotor adaptation to a visual perturbation. **a.** Experimental setup for the visual clamp disparity (VCD) task used to probe sensorimotor adaptation. During the adaptation phase, participants performed center-out reaching movements while visual feedback of the cursor was clamped at a fixed 30° angular offset from the target, independent of the actual hand trajectory. Participants were instructed to ignore the cursor and aim directly toward the target. **b.** Schematic of the task structure and movement phases. Reaches were performed across sequential cycles including no-feedback (*trials 1-10*), veridical feedback (*trials 11-20*), clamped feedback (*trials 21-100*), and a no-feedback (*trials 101-110*) phase used to assess aftereffects. Early (*trials 21-31*) and late (*trials 81-100*) epochs during clamped feedback quantify the progression of implicit adaptation. Each cycle consisted of 10 consecutive trials. **c.** Example adaptation curves for participants SCS07 (left) and SCS08 (right). Hand angle gradually shifted opposite to the direction of the clamped cursor during the adaptation phase, reflecting implicit recalibration of movement direction. Stimulation increased the magnitude of adaptation compared to stimulation off. Colored lines show the mean hand angle across movement cycles under stimulation off (blue) and stimulation on (yellow). Circular markers represent the mean hand angle for each epoch and error bars indicate the standard deviation across trials within each cycle. Vertical dashed lines indicate the transition between task phases (baseline, clamped feedback, and no-feedback). **d.** Quantification of hand angle during early adaptation, late adaptation, and aftereffect cycles for each participant. Both participants exhibited larger deviations during stimulation, indicating enhanced implicit adaptation. Early adaptation corresponds to the first cycle following the onset of clamped feedback, late adaptation corresponds to the final two cycles of the clamped feedback phase, and aftereffect corresponds to the no-feedback trials immediately following the adaptation phase. Each dot represents an individual trial. Box plots summarize the distribution of hand angles across trials under stimulation off (blue) and stimulation on (yellow). Boxes represent the median and interquartile range, and whiskers denote the minimum and maximum values. Statistical significance was assessed using two-sided bootstrapping (n = 10,000): *p < 0.05, **p < 0.01, ***p < 0.001.

Previous work has shown that adaptation arises because the perturbed visual feedback biases the perceived state of the limb, generating an error—a mismatch between perceived and intended movement outcomes. This perceived limb state reflects the integration of the visual cursor, predicted limb position (i.e., positioned at the target), and proprioceptive signals from the actual limb position^47^. To minimize this error, the motor system adapts implicitly in the opposite direction—shifting limb position away from the target—to realign the biased perceived state with the intended goal (Figure 3b). As posited by multisensory integration accounts of adaptation, disrupting proprioceptive feedback with SCS should down weight proprioceptive input and increase reliance on vision. This predicts enhanced adaptation, requiring a larger shift in limb position away from the target to realign perceived and desired states (Figure 3b).

Following a baseline period with veridical visual feedback, participants were exposed to clamped visual feedback and exhibited a gradual shift away from the target, in the direction opposite to the clamp, similarly to behaviors previously reported in able-bodied subjects^48^. With stimulation OFF, SCS07 adapted by 78.0° (95% CI: 73.6, 84.2) and SCS08 by 6.4° (95% CI: 4.9, 7.6). Both participants exhibited robust aftereffects—persistent deviations in reach direction after feedback removal—confirming robust sensorimotor adaptation to the visual perturbation.

Strikingly, with stimulation ON, sensorimotor adaptation was markedly enhanced in both participants (Figure 3d): SCS07 exhibited a 40% increase and SCS08 an 105% increase in total adaptation. These findings demonstrate that disrupting proprioceptive input enhanced implicit adaptation, consistent with increased reliance on visual feedback when proprioceptive processing is degraded. Together, these findings provide causal evidence that proprioceptive disruption induced by SCS propagate to downstream motor function and modulates sensorimotor adaptation.

### Spinal cord stimulation disrupts rapid feedback corrections

Everyday actions are routinely perturbed (e.g., recovering from a jolt while carrying a cup of coffee), requiring fast, spatially precise corrections that rely primarily on proprioceptive feedback, as visual feedback is too slow to support these responses^49,50^. As a second test of whether SCS-induced disruption of proprioception propagates to downstream motor function, we examined its impact on rapid feedback corrections.

To probe this, participants performed the arm posture perturbation (APP) task with and without stimulation (Figure 4a)^50^. Specifically, participants were tasked to maintain their impaired arm in a fixed target position while the robot applied unexpected forces, creating a perturbation in random directions, and were instructed to return to the target as quickly as possible. With vision removed, proprioception served as the primary sensory drive for these corrections. If SCS-induced proprioceptive disruptions propagate downstream, we would expect degraded feedback responses. Specifically, we would expect stimulation to result in higher postural instability, as participants would be unable to maintain their initial hand position, increased endpoint error, as participant would have difficulty returning to the target following the corrective response, and greater maximum displacement. The speed of corrective responses (deceleration time and return time), instead, will probably not be affected. Indeed, previous observations in non-human primates^51^, showed that deactivation of parietal area 5, a higher-order somatosensory region critical for proprioceptive state estimation, selectively reduced spatial accuracy (postural stability, endpoint error, and maximum displacement) while preserving response timing.

**Fig. 4.**
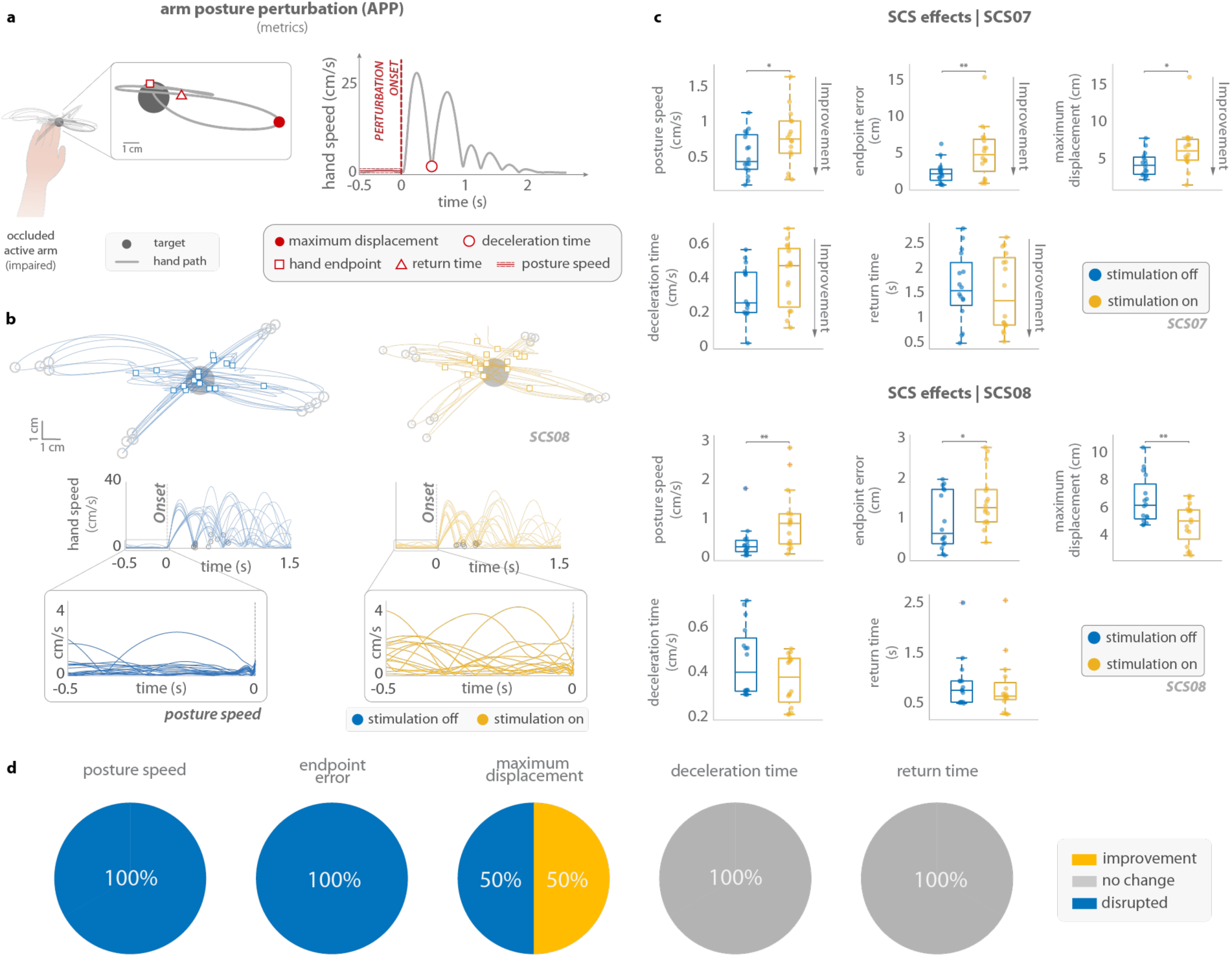
| Spinal cord stimulation disrupts rapid feedback control. **a.** Experimental setup and kinematic metrics for the arm posture perturbation (APP) task. Participants maintained their impaired arm at a central target while unexpected mechanical perturbations displaced the limb. Hand trajectories and velocity profiles illustrate the spatial metrics (endpoint error and maximum displacement) and temporal metrics (deceleration time and return time), as well as posture speed measured immediately prior to perturbation onset. **b.** Representative hand trajectories and velocity profiles from participant SCS08 during stimulation off (blue) and stimulation on (yellow). Stimulation increased baseline posture speed and produced larger endpoint errors following perturbations. **c.** Target-level comparisons of APP performance for participants SCS07 (top) and SCS08 (bottom). Box plots show the effects of stimulation on posture speed, endpoint error, maximum displacement, deceleration time, and return time across trials under stimulation off (blue) and stimulation on (yellow). Spatial control metrics were generally degraded under stimulation, whereas temporal metrics were largely preserved. Improvement was defined according to the expected direction of better performance for each metric, where reduced error in posture speed, endpoint error, maximum displacement, deceleration time and return time indicates improvement. Box plots represent the median, 25th and 75th percentiles, and minimum and maximum data points. Individual dots represent single trials. Statistical significance was assessed using two-sided bootstrapping (n = 10,000): *p < 0.05, **p < 0.01, ***p < 0.001. **d.** Summary across participants for the different APP metrics. Posture speed and endpoint error worsened in all participants during stimulation. Maximum displacement showed mixed effects, while deceleration time and return time were unchanged.

As hypothesized, when turned ON, SCS impaired postural stability and spatial control across participants (Figure 4c, Figure 4d). Indeed, posture speed increased substantially (SCS07: +78%; SCS08: +194%), and endpoint error rose in parallel (SCS07: +144%; SCS08: +53%), consistent with reduced spatial precision and increased limb instability. Maximum displacement exhibited participant-specific effects, increasing in SCS07 (+50%) but slightly decreasing in SCS08 (–18%). In contrast, temporal measures of deceleration and return time did not differ between stimulation ON and OFF, indicating that the timing of corrective responses was preserved despite degraded spatial performance (Figure 4e).

In summary, SCS induces a marked degradation in feedback control as suggested by an increase in postural stability and endpoint error, but not in the motor responses themselves.

### Spinal cord stimulation enhances goal-directed reaching

These experiments demonstrated that SCS altered kinesthesia and disrupted functional use of proprioception during implicit adaptation and rapid corrections. We then asked whether goal-oriented reaching movements would be disrupted or unaltered. For this, we employed the visually guided reaching (VGR) task (Figure 5b), which engages voluntary, goal-directed movements under continuous visual feedback. Specifically, participants performed center-out planar reaching movements from a central position toward visually displayed targets using the KINARM exoskeleton (Figure 5a). Participants were instructed to reach each target in a single, continuous movement at a rapid pace while maintaining accuracy. Importantly, stimulation did not induce consistent changes in movement speed. Indeed, mean speed, peak speed, and movement time exhibited heterogeneous changes over participants (Supplementary Figure 4a). This suggests that any observed changes in reaching performance were not attributable to a generalized increase or decrease in movement speed with stimulation ON.

**Fig. 5.**
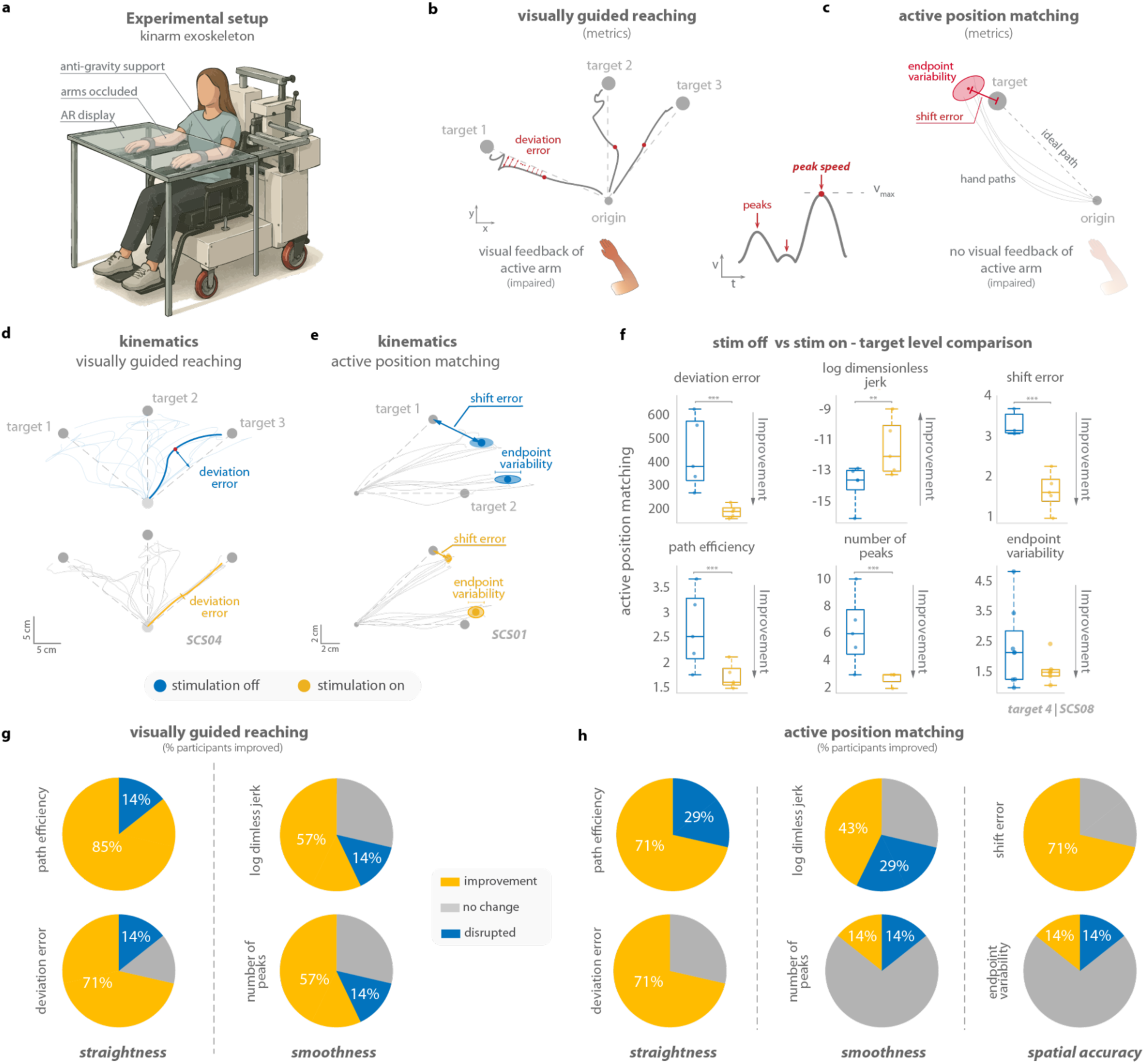
| Spinal cord stimulation enhances goal-directed reaching. **a.** Experimental setup: participants performed planar reaching using the KINARM exoskeleton using the impaired limb during the stimulation on and off conditions. **b.** Schematic of the VGR task: participants reached from a central origin to visual targets with continuous visual feedback of hand position. Movement straightness was assessed via deviation error and path efficiency; movement smoothness was quantified by velocity peaks and log-dimensionless jerk. **c.** Schematic of the APM task: participants performed similar reaches without visual feedback, using only proprioception to guide movement. In addition to straightness and smoothness, we also measured spatial accuracy (shift error) and precision (endpoint variability). **d.** Representative hand paths for VGR for participant SCS04 showing straighter and more consistent trajectories under stimulation. **e.** Representative hand paths for APM for participant SCS01 highlighting reduced shift error and endpoint variability. **f.** Box plots show target-level effects of stimulation (stimulation OFF: blue vs stimulation ON: yellow) across metrics for the APM for SCS08. Significant improvement is seen across the straightness (deviation error, path efficiency) and smoothness (log-dimensionless jerk, velocity peaks) metrics. Additionally, significant improvement was observed in spatial accuracy (shift error) without vision. Improvement was defined according to the expected direction of better performance for each metric, with reduced error (deviation error, path efficiency, number of peaks, shift error, endpoint variability) and increased and log-dimensionless jerk indicating improvement. Box plots represent the median, 25th and 75th percentiles and minimum and maximum data points. Individual dots represent single trials. Statistical significance was assessed with two-sided bootstrapping (n = 10 000): ^∗^*p* < 0.05, ^∗∗^*p* < 0.01, ^∗∗∗^*p* < 0.001. **g.** VGR pie charts summarizing the stimulation effects by metric across participants: yellow indicates improvement, blue indicates disruption, and gray indicates no change under stimulation. Results show that stimulation enhanced straightness and smoothness in most participants. **h.** APM pie charts summarizing the stimulation effects by metric across participants. Straightness metrics consistently improved while smoothness metrics showed lesser improvement. Spatial accuracy (shift error) substantially improved but not spatial precision (endpoint variability).

We analyzed kinematic features reflecting movement straightness (deviation error and path efficiency) and movement smoothness (log-dimensionless jerk and number of velocity peaks). Improved motor control would be reflected in increased movement straightness and smoothness.

Interestingly, SCS significantly improved volitional motor control across participants (Figure 5d, Supplementary Figure 3a). Specifically, stimulation improved movement straightness as quantified by path efficiency (85% of participants) as well as deviation error (71% of participants). Importantly, smoothness was also improved in most of the participants (57%) both for log-dimensionless jerk and number of velocity peaks (Figure 5g). Reaching movements primarily involved elbow-dominant or mixed shoulder-elbow coordination across targets. Importantly, stimulation did not alter joint contribution, as the reaching strategy involved similar patterns between stimulation conditions (Supplementary Figure 3a).

Overall, these results demonstrated that SCS enhances voluntary, goal-directed motor control when visual feedback is available.

We then examined movements performed without visual feedback using the Active Position Matching (APM)^52^. As in the VGR task, participants were instructed to reach each target in a single, continuous movement at a rapid pace while maintaining accuracy, but without any visual feedback of the limb throughout the movement (Figure 5c). Similar to the VGR task, stimulation did not induce a uniform increase in movement speed, with mean speed, peak speed, and movement time exhibiting heterogeneous changes across participants (Supplementary Figure 4b).

Similarly to the visually guided reaching results, SCS enhanced trajectory straightness in most participants (> 71% of them) for all the assessed metrics (Figure 5h). Improvements in movement smoothness were less consistent across participants, but no disruption was reported when the stimulation was ON suggesting that, in both conditions, participants required more correction due to the absence of vision. Importantly, shift error significantly improved with the stimulation ON in most of the participants (> 71% of them) further demonstrating an enhanced spatial accuracy (Figure 5h, Supplementary figure 3b). Notably, endpoint variability — a measure of precision — was largely unaffected by stimulation.

Together, these findings demonstrate that SCS enhanced goal-directed reaching even when visual feedback is absent and despite the disrupted proprioception.

## Discussion

Here, we leveraged cervical SCS and a comprehensive battery of upper-limb sensorimotor assays to provide a causal test of proprioception’s contribution to human motor control. We found that SCS selectively disrupts dynamic, kinesthetic proprioception while sparing static position sense. As a downstream consequence, SCS impairs online corrective responses to mechanical perturbations and amplifies sensorimotor adaptation in response to visual feedback, effectively downweighing proprioceptive information. Strikingly, ballistic, goal-oriented reaching improved with SCS ON. Together, these results provide causal evidence that proprioception is critical for online corrections yet not necessary for rapid, goal-directed actions.

### A comprehensive, causal investigation of proprioception in upper-limb motor control

For decades, the field has been divided between two views of proprioception’s role in voluntary movement. One holds that proprioception is fundamental to all voluntary movement, continuously informing the brain of the limb’s state and enabling accurate movement execution^10,12,45,53,54^. The other holds that proprioception is not necessary for fast, goal-directed reaching, pointing to deafferented animals and patients who can still produce accurate, ballistic movements toward visible targets^1–5,55^. Our results are more consistent with the latter view.

Indeed, sensorimotor behaviors that depend on proprioception for online control and recalibration were markedly affected by SCS. On the arm perturbation task, stimulation impaired the spatial accuracy and postural stability of rapid corrective responses, mirroring non-human primate findings in which parietal area 5 deactivation selectively reduces spatial accuracy while preserving response timing^51^. We also observed altered performance during implicit adaptation, suggesting greater visual reliance when proprioceptive input is degraded^47^. These results confirm that proprioception is necessary when the motor system must integrate sensory feedback to correct, stabilize, or recalibrate ongoing movement. In contrast, during both visually and non-visually guided reaching, SCS preserved movement straightness and smoothness, indicating that degraded proprioception does not preclude accurate reaching to a known target. This pattern suggests that, for movements that can be executed under feedforward control, optimized motor output is sufficient.

Because a single, reversible manipulation produced both effects in the same individuals, we can attribute this dissociation to proprioception itself rather than to compensation or population differences — moving the debate beyond the binary question of whether proprioception is necessary for movement, toward a more precise account of when it is. Proprioception is essential for online correction, postural stabilization, and adaptation to perturbations, but it is not required for fast, goal-directed reaching to a known target. This resolution explains why deafferented patients can still generate accurate center-out reaches under visual guidance and continue to produce reaching movements even without vision, albeit with reduced precision^9,56–58^, but cannot interact flexibly with unpredicted perturbations.

### Epidural spinal cord stimulation improves feedforward reaching while degrading feedback control

An important point regarding visually and non-visually guided reaching movements is that SCS did not only preserve movement straightness and smoothness, but it actually improved motor performance. It is therefore logical to ask: how can SCS simultaneously degrade proprioception and enhance goal-directed reaching? The answer may lie in what each behavior demands of the motor system. Fast, goal-directed reaches to a known target can be executed largely open-loop, requiring a well-formed descending motor command but minimal moment-to-moment sensory feedback. Online correction, postural stabilization, and adaptation, by contrast, require the motor system to integrate proprioceptive input as the movement unfolds. SCS strengths the first while disrupting the second.

Prior work from our group shows that SCS enhances cortico-muscular coherence and agonist-antagonist coordination, reflecting improved motor cortex-spinal coupling beyond a simple increase in spinal motor drive^59–61^. This strengthening of the descending command is sufficient to support fast, feedforward-controlled movements, even when proprioceptive feedback is degraded. Consistent with this view, SCS improved endpoint accuracy in the active position matching task even without visual feedback, indicating that a robust motor command can sustain – and even sharpen – reaching when sensory information is limited^62–65^. By strengthening descending drive, SCS enhances the feedforward control on which fast, goal-directed reaching depends, even if it degrades the feedback signals required for correction and adaptation.

### Clinical considerations

The transient sensory disruption under SCS does not imply proprioception is irrelevant for long-term recovery. In our clinical trial, participants reported clinically meaningful motor improvements persisting even with stimulation OFF, but only among those with preserved baseline (i.e., pre-treatment) sensory function^29^. This suggests proprioception may play a critical role in long-term motor recovery, even if not indispensable for immediate goal-directed movement.

SCS paradigms preserving sensory integrity while facilitating motor output may enhance long-term recovery outcomes, including for patients with baseline sensory deficits. For instance, Formento et al. showed that temporally patterned, phasic stimulation aligned with physiological afferent firing preserves proprioceptive signaling while maintaining motor excitation^34^. Such sensory-compliant approaches could minimize interference with dorsal root afferents neural activation and enhance the therapeutic effects of SCS, as previously shown in rodent models^33^.

In summary, these observations suggest that stimulation strategies which do not respect physiological sensory signaling may bias control toward motor output at the expense of sensory integrity, with potential implications for long-term rehabilitation. Combining SCS with proprioceptive training, instead, may promote adaptive sensorimotor reweighting. Future studies should examine how repeated continuous stimulation reshapes sensory weighting, corticospinal coupling, and motor learning, ultimately defining the boundary between sensorimotor short-term facilitation and recovery.

### Limitations of the study

A few methodological and interpretational considerations should be acknowledged when evaluating these findings. SCS likely increased afferent noise rather than fully abolishing proprioception, leaving participants with partial sensory input. Yet, this partial disruption combined with motor enhancement, still reveals that accurate reaching persists even when proprioceptive acuity declines. In addition, the robotic tasks isolated specific sensorimotor components under controlled conditions, removing gravitational and environmental variability. While enabling precise kinematic analysis, this limits generalization to 3D or distal dexterous movements, which likely demand greater sensory and corticospinal integrity. Additionally, we examined immediate rather than long-term effects. Longitudinal studies integrating SCS with rehabilitation are needed to assess sustained benefits. Participant-specific factors such as lesion location, integrity of spared pathways, and baseline sensory function likely contribute to the heterogeneity observed across individuals. Additionally, similarly to other studies^19,29^, target positions were different across participants because they were selected so that reaching movements were within the range of motion of each patient in the absence of stimulation. However, our within-subject design, where each participant served as their own control, mitigates these concerns.

These considerations do not undermine our core finding: across diverse validated tasks, SCS degraded proprioception and the feedback-dependent control that relies on it, yet improved fast, goal-directed reaching, consistent with a strengthening of feedforward motor output.

## Acknowledgements

We thank T. Simpson for technical and engineering support. This work was supported by the National Institutes of Health BRAIN Initiative (grant no. UG3NS123135-01A1 to M.C. and D.J.W.), as well as internal funding from the Office of the Provost (to E.C.), the Department of Neurological Surgery at the University of Pittsburgh (to M.C.), the Department of Mechanical Engineering and the Neuroscience Institute at Carnegie Mellon University (to D.J.W.), and the Department of Physical Medicine and Rehabilitation at the University of Pittsburgh (to E.P.).

## Author Contributions

M.C., D.J.W. and E.P. conceived the study. M.C., D.J.W. and E.P. secured the funding. E.P., E.C., and J.T. designed the experiments and assessments. M.P.P., D.J.W., N.V. and E.S. designed and implemented the stimulation control system. E.C. and E.P. designed and implemented the KINARM sensorimotor tasks. R.M.dF. and E.P. designed and performed the MRI-based analyses. A.B., R.M.dF. and G.F.W. implemented patient recruitment, eligibility and monitoring, and coordinated management of the study. M.C., L.E.F. and P.C.G. designed the neurosurgical approach. E.C., R.M.dF., N.V., L.B., performed the experiments. E.C. designed the statistical data analyses. E.C analyzed the data. E.C. created the figures. E.C., E.P., J.T., J.W.K and M.C, interpreted the results. E.C., E.P., J.T., and J.W.K. wrote the paper and all authors contributed to its editing.

## METHODS

All experimental procedures were approved by the University of Pittsburgh Institutional Review Board (IRB) under protocol STUDY19090210, in accordance with an abbreviated investigational device exemption. The study protocol is registered on ClinicalTrials.gov (NCT04512690). A total of eight participants were enrolled in the study; however, only seven completed the full protocol. One participant (SCS06) was withdrawn prior to implantation due to a newly diagnosed arrhythmia discovered during preoperative screening. All participants provided written informed consent in accordance with IRB-approved procedures. Participants were compensated for their time and reimbursed for travel and lodging expenses incurred during the study period.

### Inclusion Criteria

Participants were eligible if they were between 21 and 70 years of age and had experienced an ischemic or hemorrhagic stroke at least six months prior to study enrollment (Supplementary Table 1, Supplementary Figure 1a). All individuals presented with chronic hemiparesis affecting the upper limb and had a baseline Fugl-Meyer Assessment (FMA) score for upper-extremity motor function between 7 and 45, indicating moderate to severe motor impairment (Supplementary Figure 1b). Prior to participation, all candidates underwent a comprehensive medical screening to ensure study eligibility. Individuals were excluded if they had severe comorbidities, previously implanted medical devices, claustrophobia, or if they were pregnant or breastfeeding. Participants were also required to be free of anticoagulant, anti-spasticity, and anti-epileptic medications during the study period.

### Participant information

Seven individuals with chronic post-stroke upper-limb hemiparesis completed the study. Participants ranged in age from 31 to 64 years and were between 2 and 10 years post-stroke at the time of enrollment. All had sustained either ischemic or hemorrhagic strokes resulting in moderate to severe motor deficits, with upper-extremity Fugl-Meyer Assessment (FMA) scores ranging from 15 to 36 at baseline (Supplementary Figure 1b). Hemiparesis was present on the left side in five participants.

Stroke etiologies included right middle cerebral artery (MCA) ischemic or hemorrhagic infarcts, basal ganglia hemorrhage, lacunar infarct, and cavernous malformation affecting the thalamus and midbrain (Supplementary Figure 1a). Several participants had undergone prior interventions such as botulinum neurotoxin injections, tendon lengthening surgery, splinting, or physical and occupational therapy. In cases where botulinum toxin was previously used for spasticity management, injections were discontinued at least 6 weeks prior to enrollment, in accordance with study requirements.

Two participants (SCS01 and SCS05) had a history of cervical or neurosurgical procedures unrelated to the implant. All participants tolerated the procedures and stimulation sessions, and none experienced serious adverse events during the trial. Detailed clinical histories and baseline assessments for each participant were reported previously^29^.

### Study Design

The experiments reported here were conducted as part of a broader pilot clinical trial (ClinicalTrials.gov: NCT04512690) aimed at evaluating the safety and efficacy of epidural spinal cord stimulation for restoring upper-limb motor function in individuals with chronic post-stroke hemiparesis. The clinical trial follows a single-group, open-label, prospective design, with temporary implantation of clinical SCS leads for up to 29 days, after which the electrodes are explanted to minimize risk.

Following screening and baseline evaluations, participants underwent percutaneous implantation of SCS leads. Starting on post-implant day 4, they participated in up to 19 scientific sessions, scheduled five times per week, 4 hours per day, until the explantation day. During the first week sessions, stimulation parameters were systematically calibrated to identify individualized configurations that elicited consistent motor facilitation. These stimulation settings were then used throughout the remainder of the study.

The present study focuses on a subset of scientific sessions designed to investigate the sensorimotor mechanisms underlying SCS-driven improvements in upper-limb control. Specifically, we evaluated how SCS influenced different components of motor control, sensory integration and sensorimotor adaptation. These tasks included: threshold to detection of passive motion, passive position matching, arm posture perturbation, visual clamp adaptation, visually guided reaching, and active position matching. For each task, we compared immediate effects between stimulation ON and stimulation OFF conditions using within-subject design. These experiments were performed over the last two weeks of the study. The order of stimulation conditions was randomized across participants for each behavioral task.

Although the broader clinical trial included both primary (safety) and secondary (efficacy) outcomes, such as isometric strength, Modified Ashworth Scale (MAS), and Fugl-Meyer Assessment (FMA), the present report does not include these clinical endpoints. Instead, we focus on quantitative kinematic and proprioceptive metrics derived from the robotic assessments to better understand the neurophysiological impact of SCS on sensorimotor control. Full details of the trial’s primary and secondary outcomes are described in the main study publication and in the publicly available protocol^19,29^.

### Intraoperative Procedures

Two epidural spinal cord stimulation leads were implanted percutaneously into the epidural space over the cervical spinal cord using a standard fluoroscopy-guided procedure under general anesthesia. Intraoperative anterior-posterior and lateral fluoroscopic images were used to guide placement and estimate rostrocaudal coverage of the cervical segments. The goal was to position the two clinical leads to span spinal segments C3 to T1, enabling selective access to motor pools controlling both proximal and distal upper-limb muscles on the affected side. During the procedure, stimulation selectivity and correct lead placement were confirmed by generating recruitment curves at each contact using a monopolar intraoperative neuromonitoring system (Xltek Protektor, Natus Medical) with a subdermal return electrode placed contralateral to the affected arm. Stimulation pulses were delivered at 1–2 Hz while current was ramped from 0 to 5 mA. Compound muscle action potentials^37,38^ (CMAPs) were recorded bilaterally using subdermal needle electrodes across a broad set of muscles (including trapezius, deltoid heads, biceps, triceps, forearm flexors/extensors, and intrinsic hand muscles). Contralateral recordings ensured that SCS did not induce cross-over effects. After securing the leads, stimulation specificity was further evaluated using frequency-dependent suppression (i.e., stimulation at 20Hz) to confirm dorsal afferent activation rather than direct motor efferent recruitment. At study completion, the leads were removed. Full details regarding the surgical procedure can be found in Powell et al.^19^, and de Freitas et al.^29^.

### Imaging Procedures

A series of imaging modalities were used to confirm lead stability and to characterize participant-specific neuroanatomy. X-ray images were acquired 1–2 weeks post-implantation in both anterior-posterior and lateral views to verify that the leads remained stable and aligned with the intended cervical segments (Supplementary Figure 1c). To visualize stroke lesions, all participants underwent structural MRI using a 3T Prisma system (Siemens) with a 64-channel head and neck coil. High-resolution T1-weighted and FLAIR images were acquired and manually segmented using MRIcron, with lesion overlays visualized in MRIcroGL after 2 mm Gaussian smoothing (Supplementary Figure 1a).

### Recruitment Curves

To evaluate the selectivity of SCS in recruiting individual muscles, recruitment curves were generated by measuring the peak-to-peak amplitude of CMAPs as a function of stimulation amplitude. During behavioral testing, we repeated the recruitment curve procedure performed intraoperatively. We used a wireless EMG system (Delsys, Trigno) to record the CMAPs from multiple upper-limb muscles, such as trapezius, anterior deltoids, biceps, triceps, brachioradialis, and abductor pollicis brevis. Stimulation was delivered to one contact at a time at 1–2 Hz, while current was ramped from 0 to 8 mA, or until the participant reported discomfort. Monopolar stimulation was applied using a return surface electrode placed over the iliac crest contralateral to the affected arm. For each muscle, the peak-to-peak amplitude of CMAPs elicited at each stimulation amplitude was normalized to the maximum amplitude obtained across each measured trial. This process was repeated for every contact on both leads, thereby obtaining recruitment curves that described a relationship between SCS amplitude and muscle recruitment.

### Systematic procedure for SCS calibration

Stimulation parameters were individually calibrated to optimize voluntary upper-limb motor output while minimizing discomfort. First, recruitment curves were used to identify contacts that selectively activated different muscle groups (e.g., elbow flexors vs. extensors). Then, a combination of patient feedback, qualitative assessment, and quantitative performance comparisons were used to finalize the stimulation amplitude (between 0.2 mA and 8 mA) and frequencies (between 40 Hz and 100 Hz) used in behavioral assessments. Pulse-width was typically set to 200 μs. Frequency was only increased when increasing stimulation amplitude failed to produce satisfactory outcomes. After individually calibrating a set of contacts (usually 2 or 3), we combined them with a personalized configuration for each participant. Stimulation amplitudes were re-adjusted when combining multiple contacts to minimize interference between their effects. Less frequently, we also adjusted the stimulation frequency across contacts. Electrical stimulation was delivered using a biphasic constant-current stimulator (DS8R; Digitimer) connected to an 8-channel multiplexer (D188; Digitimer), with real-time control implemented in MATLAB.

### Behavioral Experiments

Sensorimotor function was evaluated using two robotic platforms: the KINARM Exoskeleton Robot (BKIN Technologies, Canada) and the HUMAC NORM dynamometer (CSMi, USA). The KINARM system allows bimanual planar arm movements with full gravity compensation and records high-resolution kinematic data from both arms (1000 Hz sampling rate). Participants were seated with their arms supported at shoulder height, and visual stimuli were projected onto a horizontal display aligned with the movement plane. A cloth was positioned above the arms to obstruct visual feedback during some of the tasks.

Given the presence of upper limb impairments in our cohort, we first evaluated each participant’s ability to perform reaching movements within the KINARM workspace. For the APM and PPM tasks, eight peripheral targets were arranged in a circular configuration around a central target. The radial distance from the center was set at either 10 cm or 15 cm depending on the participant’s comfortable reaching range. We initially attempted to align the central target with the KINARM’s standard anatomical reference position (shoulder abducted to ∼85°, elbow flexed to 90°). However, in cases where participants experienced spasticity, pain, or discomfort in this posture, we adjusted the central target to a more natural position tailored to their specific limitations. Peripheral targets were then positioned relative to this new center. Participants were asked to reach toward all eight directions with visual feedback, and we selected two or more directions for subsequent APM and PPM testing based on targets they could reliably reach more smoothly under visual guidance^52^.

Additionally, for the Threshold to Detection of Passive Motion (TTDPM) task, we used the HUMAC NORM, a robotic torque dynamometer designed for precise control of joint motion. This platform enabled the application of slow, robot-driven displacements to the upper limb while recording joint position with high fidelity. The system allowed for passive movements at speeds as low as 0.25°/s, essential for quantifying proprioceptive detection thresholds without eliciting stretch reflexes or voluntary responses.

#### Threshold to Detection of Passive Movement

To evaluate dynamic proprioception in the impaired arm, we implemented a passive motion detection task. For this, we used the HUMAC NORM, a robotic torque dynamometer designed for precise control of joint motion. Visual and auditory input were minimized by using a blindfold and headphones playing continuous white noise. The tested arm was positioned at 90° of elbow flexion at the start of each trial. Following a randomized delay (1-5 seconds), the HUMAC NORM passively moved the elbow joint in either flexion or extension at a constant velocity of 0.5°/s. For participant SCS08, who had difficulty perceiving movement at this speed even with stimulation OFF, trials were instead conducted at 1°/s. Participants were instructed to press a button with their unimpaired arm as soon as they detected any motion and report the sensed direction (Figure 2b). If no response was registered before the arm reached ±15° of displacement from the starting position, the trial was terminated and recorded as a non-detection. Each direction (flexion and extension) was tested six times in randomized order. We performed this assessment in 3 participants (SCS05, SCS07 and SCS08).

For the TTDPM task, we quantified two main outcome measures: detection angle and error rate (Figure 2c). Detection angle was defined as the amount of elbow joint rotation, in degrees, at the moment the participant pressed the button indicating perceived motion. Error rate was calculated as the proportion of trials in which the participant failed to detect motion or reported the wrong movement direction. Together, these metrics provided a measure of both proprioceptive sensitivity (angular displacement) and consistency or reliability of detection (error rate) for passive motion in the impaired arm.

#### Passive Position Matching

To assess static proprioception, we used a mirror matching task which has been used before^41–43,43^ using the KINARM platform. We chose this bimanual design because it most effectively engages muscle spindle afferents and provides the highest reliability for quantifying static proprioception^66,67^. The target locations were selected based on their spatial reaching capabilities. At the start of each trial, the robot moved both arms to a central position. While there, a white fingertip cursor was shown for 500 ms to provide visual recalibration of hand position. The cursor then disappeared, and the impaired arm was passively displaced by the robot to one of the preselected target locations. After the arm stabilized, auditory and visual cues were displayed (100 – 700 ms variable delay) to indicate to the participant that they should initiate a matching movement using the unimpaired arm. Participants attempted to mirror the felt position of the passive arm as accurately as possible. Upon reaching the perceived location, the participant verbally indicated completion, and the final position was recorded. Finally, the robot moved both arms back to the central position to start a new repetition. Each target was repeated five times in randomized order.

To characterize proprioceptive performance, we computed two metrics: spatial accuracy (shift error), defined as the distance between the true mirrored position and the participant’s matched position (Figure 2e, Supplementary Figure 2a), and precision (endpoint variability), defined as the trial-to-trial dispersion of the matching arm^41^ (Figure 2e, Supplementary Figure 2b). For both measures, larger values are indicative of degraded spatial accuracy and precision.

#### Clamped Feedback

The clamp feedback task was used to assess implicit sensorimotor adaptation to a persistent visuo-proprioceptive mismatch. Two participants (SCS07 and SCS08) performed center-out reaching movements using the KINARM Exoskeleton Robot in a horizontal plane with full gravity support. A white circular cursor representing the hand position was displayed on the screen, while direct vision of the limb was occluded. At the start of each trial, participants were required to move the cursor to a central target and hold it within a start window for 500 ms. Once stabilized, a single peripheral target appeared, positioned 8 cm from the center at a fixed angle of 30° counterclockwise from the vertical axis. This specific target location was selected based on prior work showing that placing targets at an angular deviation from cardinal directions (e.g. 30° counterclockwise from the vertical axis) elicits stronger implicit adaptation responses. Participants were instructed to move rapidly and directly through the target. After the radial distance was reached, the arm was returned to the central target position. The experiment consisted of four sequential blocks: (1) 10 no-feedback trials to establish baseline performance, (2) 10 veridical feedback trials to familiarize the participant with the task structure, (3) 80 clamped feedback trials to induce adaptation, and (4) 10 no-feedback trials to measure aftereffects. During veridical feedback trials, the cursor accurately tracked hand position and was extinguished when the hand crossed the 8 cm radial distance. In clamped feedback trials, the cursor trajectory was angularly offset by 30° counterclockwise relative to the target direction, moving along a fixed path regardless of the actual hand trajectory. Participants were instructed to ignore the cursor and continue aiming directly at the target. As in the veridical condition, the cursor disappeared once the hand reached the target distance. During no-feedback trials, the cursor was extinguished immediately upon movement onset and remained off throughout the reach. The task was repeated with stimulation ON and OFF, but the two conditions were tested on separate days to prevent potential carryover effects as previously reported^68^, with session order counterbalanced across participants.

The primary dependent variable for assessing implicit adaptation was movement angle, defined as the angular direction of the hand at the point where movement amplitude reached 8 cm from the start position. Movement angle quantified the angular deviation between the straight line to the target and the direction of the hand at reached amplitude. Movement angles left to the target were assigned a negative value, while those to the right of the target were assigned a positive value. Participants were instructed to move directly through the target and avoid correcting based on cursor position, ensuring that the measured movement angles primarily reflected feedforward motor planning. Adaptation was quantified by tracking changes in movement angle across clamped trials, while aftereffects were evaluated using movement angle during the final no-feedback block.

#### Arm Posture Perturbation

The Arm Posture Perturbation task was used to assess participants’ ability to respond to unexpected mechanical disturbances while maintaining a static arm posture31,32. Participants were instructed to keep their hand within a displayed target and to return to the target as quickly as possible if perturbed out of it. Once the hand cursor was placed within the target, a background torque was applied either at the shoulder (±1.5 Nm) or elbow (±1.0 Nm) over a 1000 ms period. After a random hold period of 1750–2250 ms, the load was abruptly removed over a 10 ms ramp-down, creating a mechanical “unload” perturbation. Participants then had 3 seconds to stabilize the hand back within the target zone before the trial ended. The target position corresponded to the central target from the other tasks. Visual feedback of hand position (via a white 0.5 cm fingertip cursor) was provided during an initial practice block of 8 trials. For all remaining trials, the hand cursor was removed at the moment of perturbation onset and only these trials were used for analysis. The arm and hand were occluded throughout the task using a cloth barrier to eliminate direct visual feedback. Participants completed 8 trials in the impaired arm (2 trials per condition across 4 conditions: shoulder flexion, shoulder extension, elbow flexion, and elbow extension). Trial order was randomized within blocks, and stimulation conditions were tested sequentially. We performed this assessment in 2 participants (SCS07 and SCS08).

For the APP task, we quantified five kinematic metrics to characterize participants’ responses to unexpected arm perturbations. Posture speed was defined as the peak hand velocity occurring immediately after the background load was released, capturing the intensity of the involuntary motion induced by the perturbation. Deceleration time measured the time of the first hand speed minima after forces ramp off, reflecting how quickly the limb slowed after the perturbation. Return time was defined as the time from perturbation onset to the moment the hand re-entered within 1 cm of the target, indicating how quickly participants were able to correct and stabilize their limb. Final displacement captures the distance between the hand and target center at the end of the movement, providing a measure of residual error. Maximum displacement quantified the furthest distance the hand deviated from the target during the perturbation response.

#### Visually Guided Reaching

We assessed the effects of SCS on arm dexterity using the visually guided reaching task on all participants. Each trial began with the participant’s arm passively positioned at a central starting location and locked in place. A peripheral target then appeared, and following a randomized delay of 100 – 700 ms, an auditory cue signaled the release of the arm. Participants were instructed to reach as directly and smoothly as possible toward the target. A trial was considered successful if the hand remained within a 0.5 cm radius of the target center for 500 ms. If the target was not reached within the allowed time window (10–15 s depending on the participant), the trial was terminated and the next trial began. The peripheral target locations were customized based on the participant’s available range of motion in the absence of stimulation, as done previously in similar studies^19,29^, and each target was presented six times in a randomizer order. We performed this assessment in all 7 participants.

To determine whether stimulation altered the joint coordination strategy used to perform the reach, we quantified the relative contribution of elbow and shoulder motion from the cumulative angular motion generated by each joint and classified each target as elbow-dominant, shoulder-dominant, or mixed based on the predominant strategy observed across trials. Reaching movements primarily involved elbow-dominant or mixed shoulder-elbow coordination across targets. Only target 2 for SCS01 shifted from mixed strategy to elbow dominant when stimulation was used (Supplementary Figure 3 – *Bottom Left*).

To evaluate the quality of motor execution, we computed a set of metrics characterizing both movement straightness and movement smoothness. Measures of straightness included deviation error (from peak velocity to target acquisition) calculated as the mean perpendicular distance between the real trajectory and the ideal path, which quantified spatial deviation from the ideal path and path efficiency, calculated and normalized to the straight-line distance to assess overall movement efficiency. To quantify smoothness, we computed the log dimensionless jerk, a time-and amplitude-normalized measure of the third derivative of position, and the number of velocity peaks, representing sub movements in the speed profile. Together, these metrics allowed us to assess how spinal cord stimulation influenced the execution of visually guided reaching movements.

#### Active Position Matching

The active position matching task evaluated proprioception during volitional motion without visual feedback. First, the impaired limb is positioned at the central target. For 500 ms, a white dot representing the fingertip, was displayed to provide a reference for recalibration. An auditory go cue (auditory tone with a 100 – 700 ms variable delay) was presented and the visual cursor was hidden. Simultaneously, one of the selected targets appeared on the screen. The robot then released the impaired arm, and participants performed a single, self-paced reach toward the target. They were instructed to make as straight a movement as possible and to stop and verbally report when they believed the target had been reached. The final hand position was recorded at that point, and the robot returned the limb to the center. Each target was presented five times in a randomized order. We performed this assessment in all 7 participants.

Joint dominance was quantified using the same approach described for the visually guided reaching task. Across targets, reaching movements primarily involved elbow-dominant or mixed shoulder-elbow coordination. Dominant strategy was preserved across conditions for most targets, with only three targets exhibiting stimulation-dependent changes: one shifted from elbow-dominant to mixed, one from mixed to elbow-dominant, and one from mixed to shoulder-dominant (Supplementary Figure 3 – *Bottom Right*).

For this task, we quantified the same kinematic metrics used in the visually guided reaching task that captured deviations in movement straightness and smoothness. In addition, because the APM task lacked visual feedback and required reaching based solely on proprioception, we also quantified shift error, defined as the Euclidean distance between the final hand position and the target location, and endpoint variability, computed as the standard deviation of final hand positions across trials for each target. Shift error reflects proprioceptive accuracy, while endpoint variability reflects precision. Together, these metrics allowed us to assess both motor execution quality and proprioceptive acuity under feedforward control. This assessment was performed in all the participants.

### Statistics

All statistical comparisons reported in this study were performed using a bootstrap resampling method, a nonparametric approach that does not assume normality of the underlying data distributions. For each comparison, we generated 10,000 bootstrap samples by resampling with replacement from the original data and computed the difference in means between conditions for each resample. A 95% confidence interval (CI) for the difference in means was constructed by taking the 2.5th and 97.5th percentiles of the resulting bootstrap distribution. If the CI contained zero, the result was considered not statistically significant; otherwise, the difference was considered significant. This approach provides a robust estimate of uncertainty, especially well-suited for the small sample sizes and potential distributional asymmetries typical in clinical studies. When multiple comparisons were performed simultaneously (e.g., across stimulation conditions or metrics), we applied a Bonferroni correction to adjust the significance threshold and control for Type I error.

To determine whether a participant improved, worsened, or showed no change with stimulation for a given metric, we used the same approach of de Freitas et al.^29^. We first computed the percentage change between the stimulation-on and stimulation-off conditions for each individual target. This normalization enabled comparisons across participants and targets by accounting for differences in baseline performance and the use of participant-specific target locations. We then identified statistically significant changes per target based on bootstrap-derived confidence intervals. A participant was classified as an improver if at least one target showed a significant positive change, or the majority of significant changes across targets were in the positive direction (i.e., stimulation-on was better). Conversely, participants were classified as worsened if at least one target showed a significant negative change, or the majority of significant changes were negative (i.e., stimulation-off showed better outcomes). If no targets showed significant changes, or if positive and negative changes were equally distributed, the participant was classified as no change.

## SUPPLEMENTARY FIGURES

**Supplementary Fig. 1.**
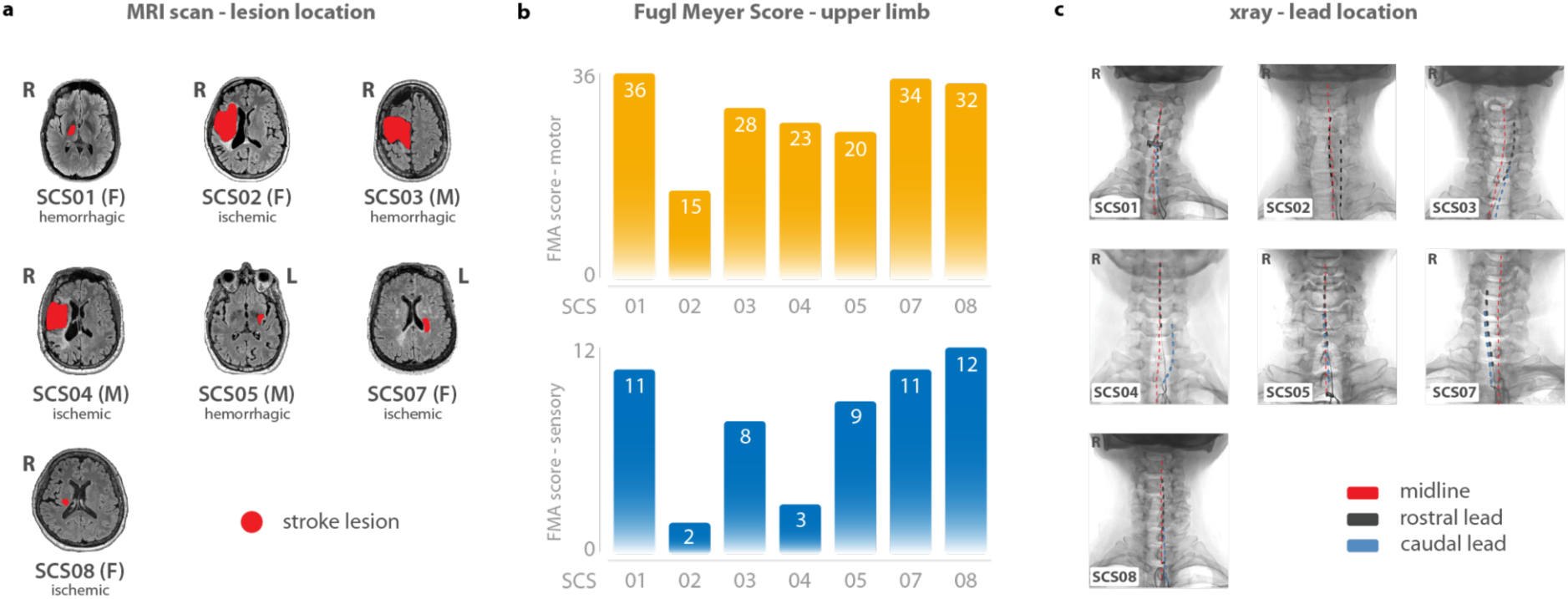
| Stroke lesion characterization, sensorimotor impairment, and electrode positioning. **a.** Axial T1-weighted MRI images from each participant showing lesion location (highlighted in red). Lesion etiology (ischemic vs. hemorrhagic) and laterality are noted; lesion sites were variable across participants. **b.** Fugl-Meyer Assessment (FMA) scores for each participant, evaluating motor (top) and sensory (bottom) function in the affected upper limb prior to stimulation. Values indicate a range of impairment, from mild to severe, across the cohort. **c.** X-ray images illustrating final epidural lead positioning relative to spinal midline for all participants. Rostral and caudal contact arrays are shown in black and blue, respectively, with red line marking midline alignment.

**Supplementary Fig. 2.**
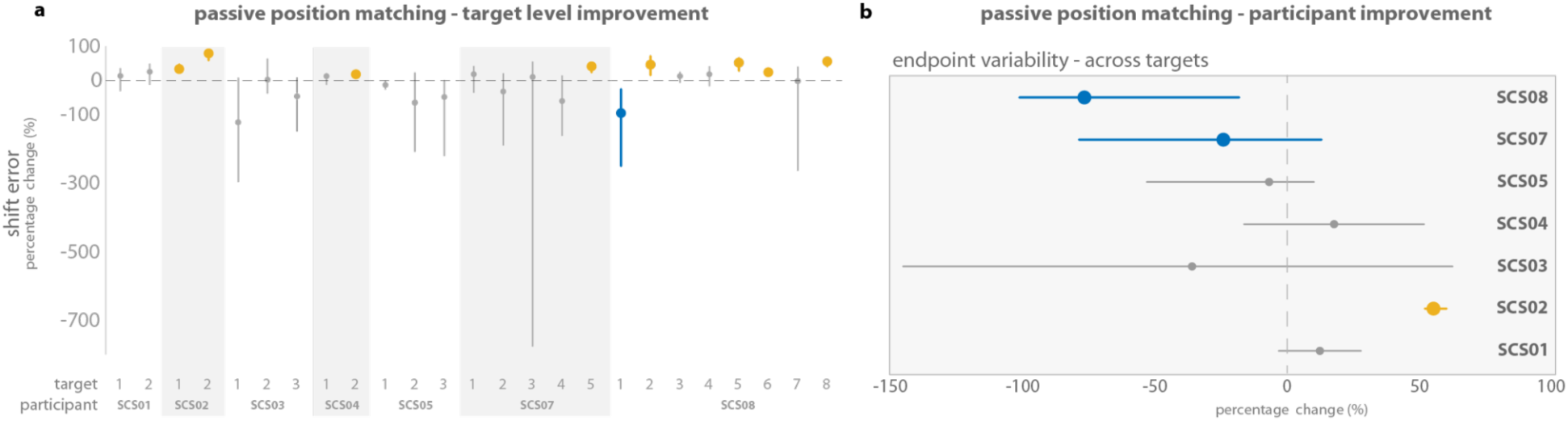
| Participant-level analysis of passive position matching performance under spinal cord stimulation. **a.** Per-target changes in shift error across all participants during the passive position matching (PPM) task. Percentage change with stimulation is shown relative to stimulation-off. Circles represent the median percentage change in shift error (accuracy): yellow indicates significant improvement (reduced error), blue indicates significant worsening (increased error), and gray denotes no change between stimulation conditions. Confidence intervals (5%–95%) of the percentage difference in shift error between SCS ON and OFF conditions are displayed with vertical lines around the median (circle). **b.** Forest plot summarizing the effect of stimulation on endpoint variability across targets for each participant. Each dot represents the median percentage change in endpoint variability (across all tested targets), and horizontal lines indicate the 95% bootstrap confidence interval. Blue indicates significant worsening (increased variability), yellow indicates significant improvement, and gray indicates no change.

**Supplementary Fig. 3.**
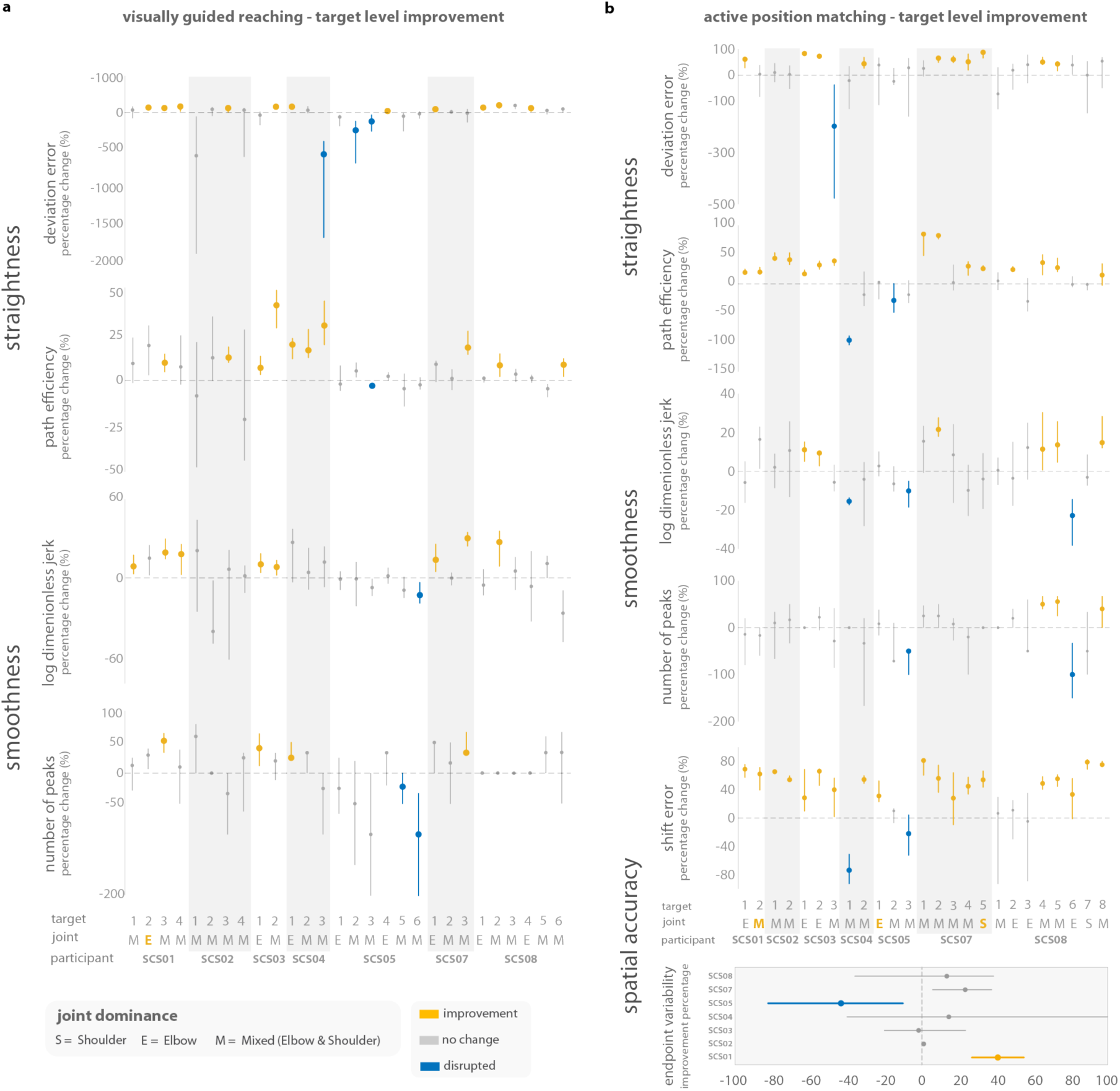
| Target-level effects of spinal cord stimulation (SCS) on trajectory kinematics in VGR and APM tasks. **a.** Visually guided reaching (VGR). Percentage change in kinematic metrics with stimulation across targets and participants, including measures of straightness (deviation error, path efficiency) and smoothness (log-dimensionless jerk, number of velocity peaks). **b.** Active position matching (APM). Same analysis applied to reaches without visual feedback, with additional spatial accuracy metrics (shift error and endpoint variability). For both plots, each dot represents the median percentage change between stimulation conditions, with vertical bars denoting 95% bootstrap confidence intervals. Yellow indicates significant improvement, blue indicates significant disruption, and gray denotes no significant change. Shaded columns group targets within participants. *Bottom inset*: Joint labels (S, shoulder-dominant; E, elbow-dominant; M, mixed shoulder-elbow) indicate the dominant joint coordination strategy for each target, determined from the relative contribution of elbow and shoulder motion. Bold labels denote targets whose dominant strategy changed with stimulation. Dominant joint usage was otherwise preserved across conditions.

**Supplementary Fig. 4.**
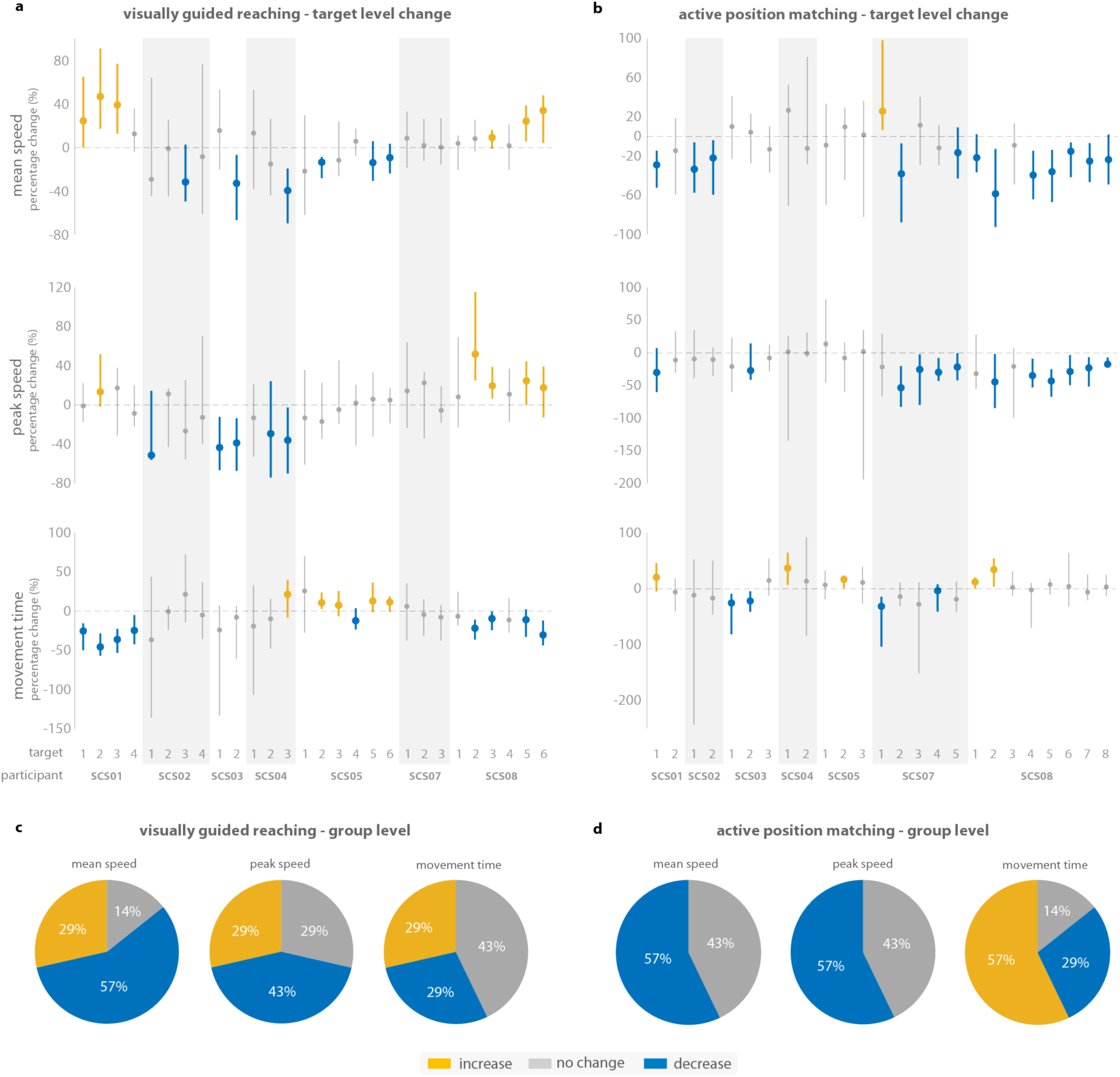
| Speed-related effects of cervical spinal cord stimulation during visually guided reaching (VGR) and active position matching (APM). **a.** Target-level percentage changes in mean speed, peak speed, and movement time under clinical stimulation relative to stimulation off are shown for each participant and target for the VGR task. **b.** Target-level percentage changes in mean speed, peak speed, and movement time under clinical stimulation relative to stimulation off are shown for each participant and target for the APM task. For panels a and b, Points represent the median percentage change, and vertical lines indicate confidence intervals. Yellow indicates significant increases under stimulation, blue indicates significant decreases, and gray indicates no significant change. **c.** Group-level pie charts summarize participant classifications based on the predominant direction of significant target-level changes for each metric (increase, decrease, or no change) for the VGR task. **d.** Group-level pie charts summarize participant classifications based on the predominant direction of significant target-level changes for each metric (increase, decrease, or no change) for the APM task. Across both tasks, speed-related metrics exhibited heterogeneous responses with no consistent stimulation-dependent increase in movement speed or decrease in movement time across participants. In contrast to the robust improvements observed in trajectory quality and smoothness metrics, these findings suggest that the effects of stimulation cannot be explained by a generalized increase in execution speed. Stimulation appears to preferentially influence movement quality and control independent of overall movement speed.

**Supplementary Table 1.** Participant demographics.

| Subject ID | SCS01 | SCS02 | SCS03 | SCS04 | SCS05 | SCS07 | SCS08 |
| --- | --- | --- | --- | --- | --- | --- | --- |
| **Sex** | F | F | M | M | M | F | F |
| **Age** | 31 | 47 | 46 | 43 | 64 | 61 | 60 |
| **Ethnicity** | caucasian | caucasian | caucasian | black | caucasian | black | asian |
| **Stroke Type** | hemorrhagic | ischemic | hemorrhagic | ischemic | hemorrhagic | ischemic | ischemic |
| **Lesion Hemisphere** | right | right | right | right | left | left | right |
| **Time after Stroke** | 9 yrs | 3 yrs | 7 yrs | 5 yrs | 2 yrs | 10 yrs | 4 yrs |
| **FM Motor (baseline)** | 36 | 15 | 28 | 23 | 20 | 34 | 32 |
| **FM Sensory (baseline)** | 11 | 2 | 8 | 3 | 9 | 11 | 12 |

**Supplementary Table 2.**
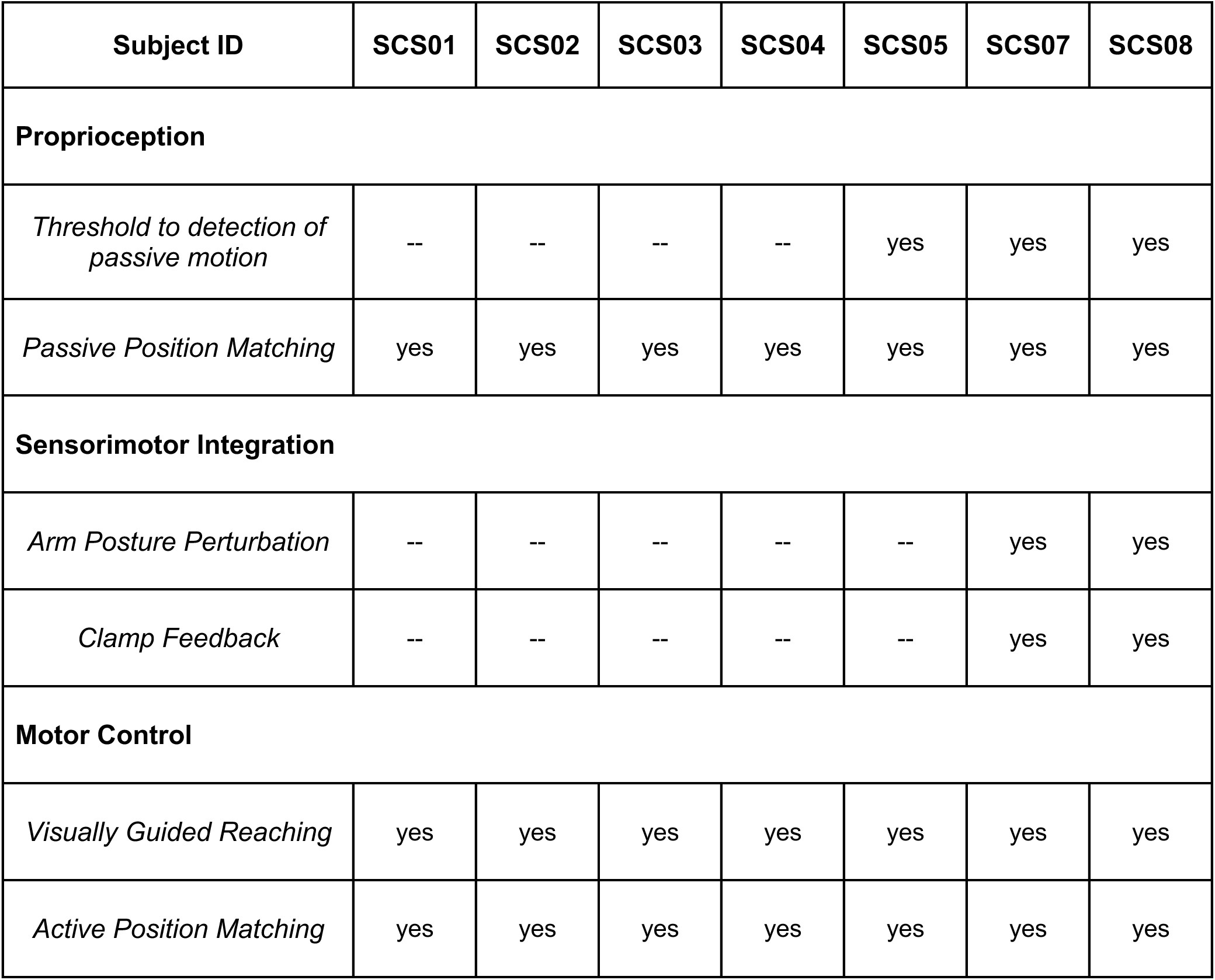
Experimental Participation.

| Subject ID | SCS01 | SCS02 | SCS03 | SCS04 | SCS05 | SCS07 | SCS08 |
| --- | --- | --- | --- | --- | --- | --- | --- |
| **Proprioception** |  |  |  |  |  |  |  |
| *Threshold to detection of passive motion* | -- | -- | -- | -- | yes | yes | yes |
| *Passive Position Matching* | yes | yes | yes | yes | yes | yes | yes |
| **Sensorimotor Integration** |  |  |  |  |  |  |  |
| *Arm Posture Perturbation* | -- | -- | -- | -- | -- | yes | yes |
| *Clamp Feedback* | -- | -- | -- | -- | -- | yes | yes |
| **Motor Control** |  |  |  |  |  |  |  |
| *Visually Guided Reaching* | yes | yes | yes | yes | yes | yes | yes |
| *Active Position Matching* | yes | yes | yes | yes | yes | yes | yes |

**Supplementary Table 2.** Fugl Meyer Sensory Score (Stimulation OFF vs ON)

| Subject ID | SCS01 | SCS02 | SCS03 | SCS04 | SCS05 | SCS07 | SCS08 |
| --- | --- | --- | --- | --- | --- | --- | --- |
| **Week 2** |  |  |  |  |  |  |  |
| *SCS OFF* | 9 | 1 | 5 | 3 | 9 | 9 | 12 |
| *SCS ON* | 9 | 1 | 5 | 1 | 10 | 9 | 12 |
| **Week 4** |  |  |  |  |  |  |  |
| *SCS OFF* | 11 | 1 | 7 | 4 | 9 | 10 | 12 |
| *SCS ON* | 11 | 1 | 7 | 1 | 9 | 12 | 12 |

